# Comparative genomics of macaques and integrated insights into genetic variation and population history

**DOI:** 10.1101/2024.04.07.588379

**Authors:** Shilong Zhang, Ning Xu, Lianting Fu, Xiangyu Yang, Yamei Li, Zikun Yang, Yu Feng, Kaiyue Ma, Xinrui Jiang, Junmin Han, Ruixing Hu, Lu Zhang, Luciana de Gennaro, Fedor Ryabov, Dan Meng, Yaoxi He, Dongya Wu, Chentao Yang, Annalisa Paparella, Yuxiang Mao, Xinyan Bian, Yong Lu, Francesca Antonacci, Mario Ventura, Valery A. Shepelev, Karen H. Miga, Ivan A. Alexandrov, Glennis A. Logsdon, Adam M. Phillippy, Bing Su, Guojie Zhang, Evan E. Eichler, Qing Lu, Yongyong Shi, Qiang Sun, Yafei Mao

## Abstract

The crab-eating macaques (*Macaca fascicularis*) and rhesus macaques (*M. mulatta*) are widely studied nonhuman primates in biomedical and evolutionary research. Despite their significance, the current understanding of the complex genomic structure in macaques and the differences between species requires substantial improvement. Here, we present a complete genome assembly of a crab-eating macaque and 20 haplotype-resolved macaque assemblies to investigate the complex regions and major genomic differences between species. Segmental duplication in macaques is ∼42% lower, while centromeres are ∼3.7 times longer than those in humans. The characterization of ∼2 Mbp fixed genetic variants and ∼240 Mbp complex loci highlights potential associations with metabolic differences between the two macaque species (e.g., *CYP2C76* and *EHBP1L1*). Additionally, hundreds of alternative splicing differences show post-transcriptional regulation divergence between these two species (e.g., *PNPO*). We also characterize 91 large-scale genomic differences between macaques and humans at a single-base-pair resolution and highlight their impact on gene regulation in primate evolution (e.g., *FOLH1* and *PIEZO2*). Finally, population genetics recapitulates macaque speciation and selective sweeps, highlighting potential genetic basis of reproduction and tail phenotype differences (e.g., *STAB1*, *SEMA3F*, and *HOXD13*). In summary, the integrated analysis of genetic variation and population genetics in macaques greatly enhances our comprehension of lineage-specific phenotypes, adaptation, and primate evolution, thereby improving their biomedical applications in human diseases.

## INTRODUCTION

Comparative and population genomics significantly improve our understanding of speciation, genetic differences, and adaptation in evolutionary biology^1–6^. Characterizing the genetic differences, especially for major alleles between two species, is the basis for revealing lineage-specific genetic changes, and their potential functions in phenotypic differences and adaptation^5,7–10^. Most comparative and population genomics studies have primarily focused on single-nucleotide variants (SNVs) or simple structural variation^9,11^. This has constrained our ability to gain insights into a broader spectrum of genetic variation (e.g., complex genomic regions)^8,10,12–14^.

With the rapid advancement of long-read sequencing techniques and hybrid assembly strategies, we have entered an era where we can generate complete telomere-to-telomere (T2T) genome assemblies^7,12,15^. These efforts enable the study of complex genomic regions, including centromeres, segmental duplications (SDs), and structurally divergent regions (SDRs, or complex regions)^10,16–20^. Currently, haplotype-resolved genome assemblies can be generated with trio-based, Hi-C-based, or Strand-seq-based approaches^21–23^. These assemblies can then be utilized to construct high-quality pangenome graphs, enhancing our understanding of genetic structure and the evolutionary history of complex genomic regions^24–26^. However, generating haplotype-resolved diploid T2T genome assemblies remains challenging, primarily due to the complexities associated with phasing centromeres, subtelomeres, and SDRs^22^.

The *Macaca* genus, including over 20 species, is one of the most adaptable and widely distributed nonhuman primates (NHPs) and it diverged from humans ∼25 million years ago (Mya)^11,27,28^. The close genetic relationship and physiological similarities shared by humans and macaques make them ideal NHP biomedical models^11,27–31^. In addition, the critical phylogenetic position of macaques (basal group of *Catarrhini*) in primate phylogeny makes them indispensable for evolutionary comparisons to shed light on lineage-specific genetic changes related to primate adaptation and human disease susceptibility^11,32–35^.

Rhesus macaques (*Macaca mulatta*, MMU) and crab-eating macaques (*M. fascicularis*, MFA) are the most used NHP models in biomedical research. MMU is frequently used in neurobiology research, yet its applicability in reproductive and developmental studies (e.g., embryogenesis) is limited due to its seasonal ovulation patterns^28,29^. In contrast, MFA exhibits year-round continuous ovulation and smaller body sizes relative to MMU^28,29^. Thus, MFA has been widespread used in research related to reproduction, development, gene editing, and drug testing in recent years^36–45^. Because of these differences, the characterization of major genetic differences between these two macaque species and between macaques and humans can further optimize the biomedical application of these two important NHP models, as well as deepen our understanding of the primate evolutionary history. Despite its importance both, the reference genomes remains largely incomplete with hundreds of gaps, collapsed regions, and incomplete gene annotations^11,27,46^. Consequently, the genetic structure of complex regions remains poorly understood in macaques. Meanwhile, the studies of large-scale genomic differences between macaques and humans are relatively limited^11,47^.

In this study, we generated a T2T haploid macaque genome assembly and 20 haplotype-resolved long-read assemblies (10 MFA and 10 MMU). Additionally, we performed long-read RNA sequencing (RNA-seq) across 15 tissues of MFA and MMU (∼30.1 million full-length cDNA transcripts). High-coverage whole-genome sequencing (WGS) was conducted for 151 macaques. With these newly generated datasets, we aim to (1) characterize the T2T macaque genome assembly and its gene annotation, (2) identify the major genetic differences between MFA and MMU with a draft pangenome, (3) analyze the large-scale genomic differences between humans and macaques, and (4) refine population history and selection in macaques.

## RESULTS

### A complete genome sequence of a crab-eating macaque

To better reconstruct a highly accurate and complete macaque reference genome assembly, we generated embryos from activated oocytes of MFA through parthenogenesis^48^. After isolation of the inner cell mass of parthenogenetic blastocysts, the genomes of the embryonic stem cells (MFA582-1) from the inner cell mass undergo self-duplication, resulting in a stable diploid state with a 42, XX karyotype that is almost entirely homozygous^48^ (“haploid” genome) (Figure 1a, Methods). Karyotype analysis and WGS were used to confirm the stability of the MFA582-1 cells and their near-uniform homozygosity (Supplementary Figures 1-3).

**Figure 1.**
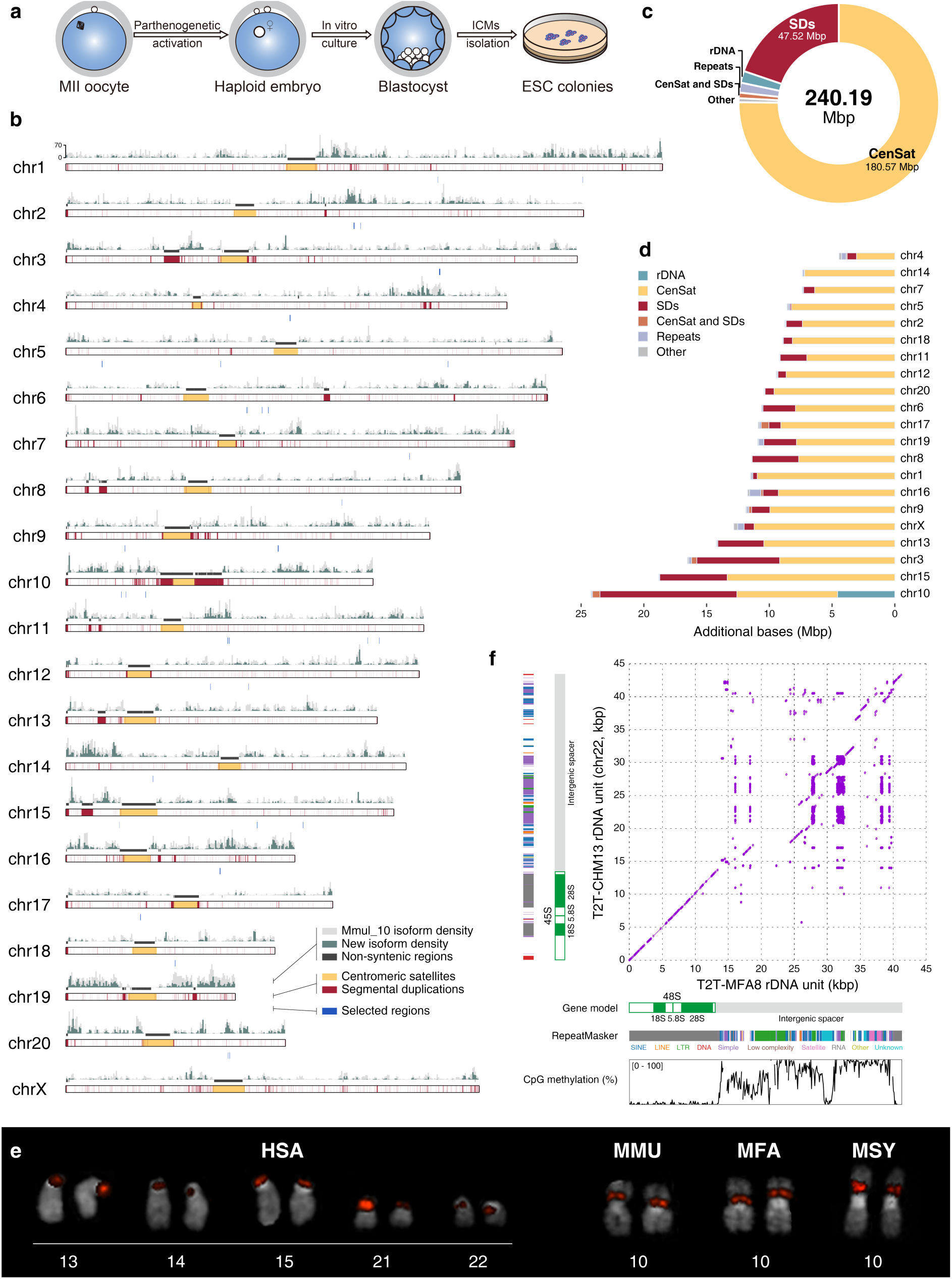
Summary of the T2T-MFA8 and rDNA validation. **(a)** Flow chart illustrating the generation of parthenogenetic embryonic stem cells (ESCs) for genome assembly. ICMs: inner cell masses. **(b)** Ideogram of T2T-MFA8 assembly with various features. The ideogram track shows the centromeric satellites (yellow) and segmental duplications (SDs; red). The gene isoform density (light gray and green), the non-syntenic regions between T2T-MFA8 and Mmul_10 (>100 kbp, dark gray), and selected regions between two species (blue) are shown above and below the ideogram track. **(c)** Pie chart illustrating the total length of added sequences. **(d)** The type of added sequences for each chromosome. **(e)** Fluorescence *in situ* hybridization (FISH) validation confirming the presence of rDNA exclusively on the chr10 of macaques. HSA: *Homo sapiens*, MMU: *Macaca mulatta*, MFA: *Macaca fascicularis*, MSY: *Macaca sylvana*. **(f)** Syntenic comparison of rDNA units between T2T-MFA8 and T2T-CHM13 (chr22). The dot plot demonstrates a conserved synteny in the rDNA coding regions between humans and macaques. The common repeat annotation and methylation pattern are listed along the axes.

We sequenced MFA582-1 with multiple sequencing technologies, including 53× PacBio high-fidelity (HiFi) sequencing with a read N50 of 19.97 kbp, 77× Oxford Nanopore (ONT) ultra-long sequencing with a read N50 of 94.68 kbp, 32× Illumina PCR-free sequencing, and 156× Illumina Hi-C library (Supplementary Figure 4 and Supplementary Table 1). Initially, we utilized HiFi and ONT reads to generate a high-quality genome assembly with a quality value (QV) of 42.28 using the haploid mode in Verkko^22^. The initial assembly shows a near-gapless configuration, with only 19 remaining gaps (147 unassigned contigs) within the assembly (Supplementary Table 2). Of these, 3 are in centromeres and 16 are in SDs.

To fill the 19 gaps presented in the initial assembly, we first used the local genomic region assembly and singly unique nucleotide *k*-mer (SUNK) mapping assembly approaches^49^, which resolved 18 complex genomic regions (Methods). Meanwhile, to decipher the remaining unresolved region mapping to rDNA, we utilized Ribotin to construct a consensus “morph”^50^ (Supplementary Figure 5). Concurrently, the *k*-mer multiplicity was employed to estimate the count of 110 rDNA copies (Supplementary Figure 6). Overall, by integrating these approaches, we successfully reconstructed a gap-free assembly. For further refinement, we manually curated 96 large structural errors and 6,629 small errors by mapping Illumina, HiFi, and ONT reads to the assembly^15,25,51^. Finally, after telomere patching and adding the rDNA array, a complete and high-quality T2T assembly of the macaque genome (T2T-MFA8v1.0) was successfully reconstructed with an NG50 of 162.13 Mbp (Figure 1b). This assembly and relative data along with a track hub are available on GitHub (https://github.com/zhang-shilong/T2T-MFA8).

### Quality assessment, genome statistics, and segmental duplication

The T2T-MFA8v1.0 assembly comprises about 3.04 Gbp of nuclear DNA and 15,561 bp of mitochondrial DNA (Table 1). This complete assembly has 240.19 Mbp additional (non-syntenic) nuclear sequences compared to the recently published Indian rhesus macaque genome assembly (Mmul_10)^11^, 231.17 Mbp to the Chinese rhesus genome assembly (rheMacS)^27^, and 267.95 Mbp to the MFA genome assembly (macFas6), respectively (Supplementary Figure 7). These newly added sequences include centromeric satellites (75.18%), non-satellite SDs (19.78%), and rDNA regions (1.90%), compared to Mmul_10 (Figure 1c and 1d).

**Table 1.** Genome statistics of T2T-MFA8v1.0 and comparison of macaque genome assemblies.

| Statistics | Mmul_10 | T2T-MFA8v1.0 | Difference (±%) |
| --- | --- | --- | --- |
| **Summary** |  |  |  |
| Assembled bases (Gbp) | 2.96 | 3.04 | +2.70 |
| Unplaced bases (Mbp) | 117.34 | 0 | -100 |
| Gap bases (Mbp) | 33.84 | 0 | -100 |
| Number of Contigs | 3,182 | 21 | -99.34 |
| Contig NG50 (Mbp) | 45.49 | 162.13 | +256.38 |
| **Gene annotation** |  |  |  |
| Number of genes | 32,528 | 33,095 | +1.74 |
| Number of protein-coding genes | 20,924 | 21,081 | +0.75 |
| Number of transcripts | 86,815 | 109,756 | +26.43 |
| Number of protein-coding transcripts | 67,292 | 92,057 | +36.80 |
| **Segmental duplications** |  |  |  |
| Percentage of segmental duplications (%) | 2.36 | 3.74 | not applicable |
| Segmental duplication bases (Mbp) | 69.90 | 113.75 | +62.73 |
| Number of segmental duplications | 33,286 | 100,829 | +202.92 |
| **RepeatMasker** |  |  |  |
| Percentage of repeats (%) | 48.56 | 54.52 | not applicable |
| Repeat bases (Mbp) | 1,437.07 | 1,659.13 | +15.45 |
| Long interspersed nuclear elements (Mbp) | 609.18 | 605.84 | -0.55 |
| Short interspersed nuclear elements (Mbp) | 399.64 | 403.89 | +1.06 |
| Long terminal repeats (Mbp) | 253.49 | 263.32 | +3.88 |
| Satellite (Mbp) | 24.78 | 200.58 | +709.54 |
| DNA (Mbp) | 102.82 | 105.20 | +2.31 |
| Simple repeat (Mbp) | 36.66 | 41.12 | +12.17 |
| Low complexity (Mbp) | 6.43 | 6.81 | +5.87 |
| Retroposon (Mbp) | 0.05 | 0.21 | +301.00 |
| rRNA (Mbp) | 0.19 | 1.58 | +720.89 |

The quality of the T2T-MFA8v1.0 assembly was validated with the same approaches used for T2T-CHM13 and T2T-CN1^15,25^ (Methods). The sizes of the assembled rDNA regions (∼4.56 Mbp) match the estimation obtained from the ddPCR experiments (Supplementary Figure 8), and the chromosome topological structures from the Hi-C interaction map show no sign of misassembly (Supplementary Figure 9). In addition, we performed quality assessments on T2T-MFA8v1.0 using Merqury^52^, compleasm^53^, and Flagger^26^. The results showed that the consensus accuracy of the T2T-MFA8v1.0 assembly equates to approximately one error *k*-mer per 10 Mbp (QV=71.44) and 99.56% genome completeness, with a potential 6.78 Mbp of incorrectly assembled bases (0.22% of the total assembly length) (Supplementary Figure 10 and Supplementary Table 3). All these genomic statistics and quality assessment indices were comparable to the current gold standard human genome assembly, T2T-CHM13^15^. Interestingly, in contrast to human genomes, there is only one rDNA region in the macaque genome^15^. We used fluorescence *in situ* hybridization (FISH) to validate this observation in different macaque species, including MFA, MMU, and a barbary macaque (*Macaca sylvanus*, MSY) (Figure 1e). The coding region within the rDNA is highly conserved between T2T-MFA8v1.0 (unit length: ∼41 kbp) and T2T-CHM13 (unit length: ∼43 kbp) and exhibits whole-gene hypomethylation (Figure 1f and Supplementary Figure 11). However, two repeats (LTR/ERV1 and α-satellite) show lineage-specific presence in the intergenic spacer of T2T-MFA8v1.0 compared to T2T-CHM13 (Figure 1f).

In the previous macaque assembly (Mmul_10), 62.73% of the SDs (≥ 1 kbp and 90% sequence identity) are unresolved or unlocalized^11^ (Supplementary Table 4); here, the T2T-MFA8v1.0 assembly achieves a full resolution of 113.75 Mbp of SDs (3.74% of the genome). Of these, 56.12 Mbp are interchromosomal SDs, and 100.47 Mbp are intrachromosomal SDs (Supplementary Figures 12 and 13). This significant advancement allows us to address a longstanding question regarding the differences in the content and genome organization of SDs between macaques and humans^19,54–56^ (Supplementary Table 5). Compared to the human genome^19^ (T2T-CHM13), the length of SDs in the macaque genome (T2T-MFA8v1.0, 113.75 Mbp, no chrY) is 78 Mbp less than that in the human genome (T2T-CHM13v2.0, 192 Mbp, no chrY). Meanwhile, the macaque genome exhibits a higher tendency for SDs to be represented as intrachromosomal states when considering both the length of SDs (chi-square test, *P* < 2.2×10^-16^) and the number of SD pairs (chi-square test, *P* < 2.2×10^-16^) (Supplementary Tables 6 and 7). Human SDs are 10 times more enriched in pericentric regions than macaque SDs (chi-square test, *P* < 2.2×10^-16^), while macaque SDs are 2 times more enriched in subtelomeric regions (chi-square test, *P* < 2.2×10^-16^) (Supplementary Tables 8 and 9).

### Gene prediction, novel genes, and gene fusion

We deeply sequenced ∼15.27 million full-length non-chimeric transcripts (FLNCs) of MFA with a long-read technology (Iso-Seq) across 15 tissues, including brain, lung, kidney, and others (Supplementary Table 10). We predicted 21,081 protein-coding genes (92,057 transcripts) and 12,014 noncoding genes (17,699 transcripts) in T2T-MFA8v1.0 with two independent approaches (Table 1; Methods). Compared to the gene annotation of Mmul_10, we added 25,113 transcripts in the previously annotated genes in Mmul_10 and found 189 novel protein-coding genes in T2T-MFA8v1.0, including 77 previously unannotated genes and 112 fusion genes (Figure 2a and Supplementary Figure 14). Of these, some genes are functionally important (e.g., *SLC25A15*), but they were absent from the previous macaque genome assembly (Mmul_10) (Figure 2b). *SLC25A15* was unannotated due to a gap in the Mmul_10 assembly. *SLC25A15* encodes the mitochondrial ornithine transporter 1 (ORNT1 or ORC1) and plays a role in serine transportation^57^. Pathogenic mutations in *SLC25A15* are associated with Hyperornithinemia-Hyperammonemia-Homocitrullinuria syndrome in humans^58,59^. Therefore, the incomplete gene annotation in the previous macaque reference genome could potentially impact the biomedical applications of macaques.

**Figure 2.**
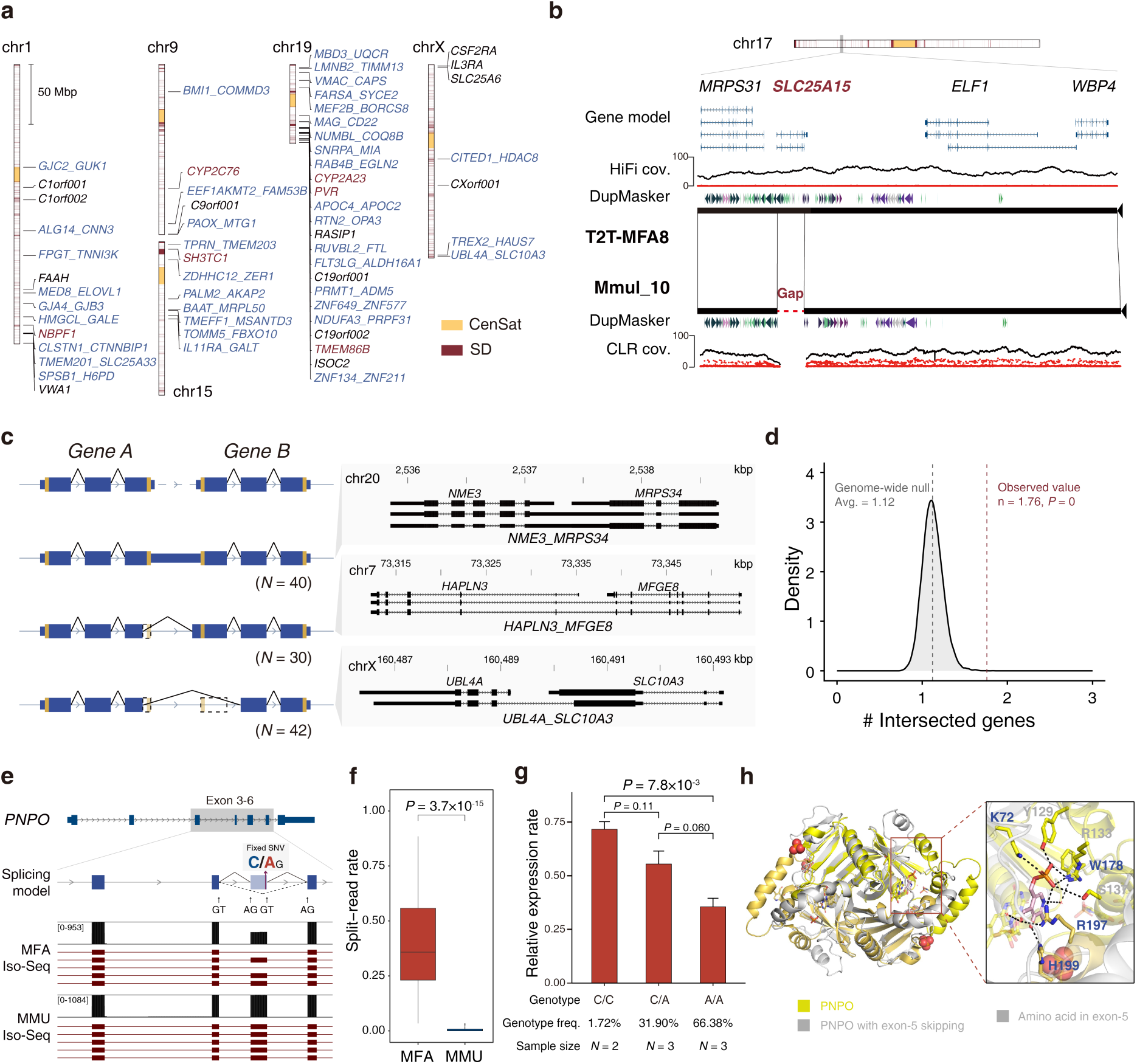
Novel gene, isoform, and alternative splice site. **(a)** Ideogram of five chromosomes with novel genes. Newly identified fusion genes are shown in blue, while other genes in segmental duplications (SDs) and other genomic regions are shown in red and black, respectively. Genes unable to be found in NCBI are marked with “CXorfXXX”. **(b)** Example of newly identified genes. The syntenic relationship between T2T-MFA8 and Mmul_10 is shown with minimiro. The red dashed line represents a 21 kbp unassembled region in Mmul_10. Gene models are shown on the top with read-depth validation below. CLR: continuous long reads. **(c)** Schematic illustration of gene fusion. Three types of gene fusion are shown on the left panel. The right panel shows examples of gene fusion events (stop codon skip, and start & stop codon skip) with support from full-length isoforms. **(d)** Gene fusion in a high gene density region. The gray represents the distribution of the number of genes adjacent to a random gene at the genome-wide level (null distribution), while the red line represents the observation of the number of genes adjacent to a fusion gene (permutation test, empirical *P* = 0). **(e)** Example of splice site alteration between MFA and MMU. The gene model of *PNPO* is presented on the top, with long-read full-length RNA-seq reads from each macaque displayed below. A fixed genetic variant between MMU and MFA is also shown (CG→AG) within the gene models. **(f)** The short-read RNA-seq confirms the exon skipping in MFA. The y-axis refers to the split-read rate of exon-5 of *PNPO*. **(g)** The qPCR validation supports that the genotypes (C/C, C/A, and A/A) are associated with exon-5 skipping in MFA. The sample size and genotype frequencies are listed below. **(h)** The predicted protein structures of PNPO with and without exon-5 show the potential loss of enzyme activity due to disrupted interactions. The right panel shows the amino acids (K72, Y129, R133, S137, W178, R197, H199) in the active center. Y129, R133, and S137 marked with gray are in the exon-5 of *PNPO*.

Full-length RNA-seq has enabled the identification of previously overlooked fusion genes (also known as chimeric RNA) resulting from transcriptional readthrough or intergenic splicing^60–62^. Fusion genes have been studied in plants and human cancer cells, but previous studies argue their presence in human normal physiology as well^60–64^. Here, we observed 112 fusion genes expressed in healthy macaques (FL counts ≥ 5) and validated them with short-read RNA-seq data from different MFA individuals^65^ (Figure 2c, Supplementary Tables 11 and 12, and Supplementary Figure 15). Of particular interest is the potential amino acids changes in the *UBL4A_SLC10A3* fusion gene. The predicted protein structure showed a spatial shift in alpha helix 9 of *SLC10A3* due to alterations in interactions and hydrophobic forces caused by the depletion of three alpha helices, relative to the two progenitor genes^66^ (Supplementary Figure 16). In addition, we found that the number of genes located in the 10 kbp flanking regions of the fusion genes is 1.57 times higher than that of the null distribution (permutation test, empirical *P* = 0) (Figure 2d, Methods). This result suggests that gene fusion is more likely to occur in genomic regions with high gene density.

### Comparative long-read transcriptomics of macaques

We also deeply sequenced 14.82 million FLNCs of MMU with Iso-Seq across the corresponding 15 tissues (Supplementary Table 10) and used these data to examine the isoform-level alternative splicing differences between MFA and MMU. We identified 795 alternative splicing differences spanning 577 genes (e.g., *PNPO*, *TBC1D8B*, and *AXIN2*) (Supplementary Figure 17 and Supplementary Table 13). These genes are enriched in DNA repair (*P* = 3.68×10^-4^), mitotic G2/M transition checkpoint (*P* = 1.23×10^-3^), Zinc transport (*P* = 4.88×10^-3^), and more^67^ (Supplementary Table 14). Among 209 coding exon skipping events, we validated 110 exon skipping differences (validation rate: 52.63%) between MFA and MMU using the previous short-read RNA-seq data^65^ (Methods). Specifically, 12 high-confidence exon skipping events exhibited at least a 20% difference in the degree of exon skipping between the two species (Methods).

Of particular interest, we observed that exon-5 of *PNPO* is alternatively skipped in all tissues of MFA, but not in those of MMU (Figure 2e). This observation was further confirmed through short-read quantification and PCR validation (Figure 2f and Supplementary Figure 18). Notably, a variant with different allele frequencies (AF) between MMU and MFA were identified in the *PNPO* exon-5 (AF for C allele in MMU: 99.4%; and in MFA: 17.7%; AF for A allele in MMU: 0.6%; and in MFA: 82.3%) (Figure 2e). We hypothesized that the C→A alteration likely introduces a novel canonical splice acceptor site in *PNPO*, potentially contributing to the observed exon-5 skipping. Subsequently, we assessed the relative expression of exon-5 and exon-3 of *PNPO* by RT-qPCR in MFA individuals with the A/A, C/A, and C/C genotypes. Our findings showed significantly reduced exon-5 expression in individuals with A/A genotypes compared to those with C/C genotypes (Figure 2g). This result supports that the exon skipping event of *PNPO* is likely related to the C/A difference (T2T-MFA8, chr16:58,789,504-58,789,504). *PNPO* is a critical enzyme involved in the vitamin B6 metabolism pathway, and mutations in *PNPO* may result in epileptic encephalopathy in humans^68,69^. The predicted protein structure of PNPO without exon-5 shows a disruption of the active site^66^ (Figure 2h). However, it remains uncertain whether the A/A genotype in MFA would lead to epilepsy or not depending on factors such as diet preference, endogenous physiological compensation, or other mechanisms, and thus further testing is necessary to understand the functions of different *PNPO* isoforms in MFA. It is noteworthy that the C/C genotype is prevalent at the orthologous site in humans^70^ (GRCh38, chr17:47,945,984-47,945,984; AF: 0.99999, gnomAD v4.0).

### Genetic differences and a draft pangenome graph

To better understand the major genetic differences between MFA and MMU, we sequenced five genetically unrelated individuals of each species with both long-read HiFi (mean coverage: 47×) and ONT (mean coverage: 27×) (Supplementary Figure 19). Utilizing this 10 individual dataset, we assembled 20 haplotype-resolved genome assemblies with trio-based (n=7) or Hi-C-based (n=3) approaches (Supplementary Table 15). The average total length of the assemblies with chromosome X or Y is approximately 3.09 Gbp or 2.95 Gbp, respectively (Supplementary Table 15). Notably, the average NG50 of these 20 macaque genome assemblies is 88 Mbp, markedly surpassing those of the human genome assemblies in the Human Pangenome Reference Consortium (HPRC) phase I^26^ (average NG50: 40 Mbp) (Figure 3a). The assembly accuracy assessment shows that the average QV is about 61.01^52^ (indicating 1 base error per 1 Mbp). For the trio-based assemblies, Illumina reads indicated a switch error rate of 1.00% and a Hamming error rate of 0.73% (Supplementary Figure 20). In addition, Flagger^26^ was employed to estimate the extent of misassembled regions, which indicated that each assembly contained only approximately 20 Mbp (∼0.65%) inaccuracies on average (Figure 3b and Supplementary Table 16). Therefore, the 20 haplotype-resolved macaque assemblies are highly contiguous and accurate.

**Figure 3.**
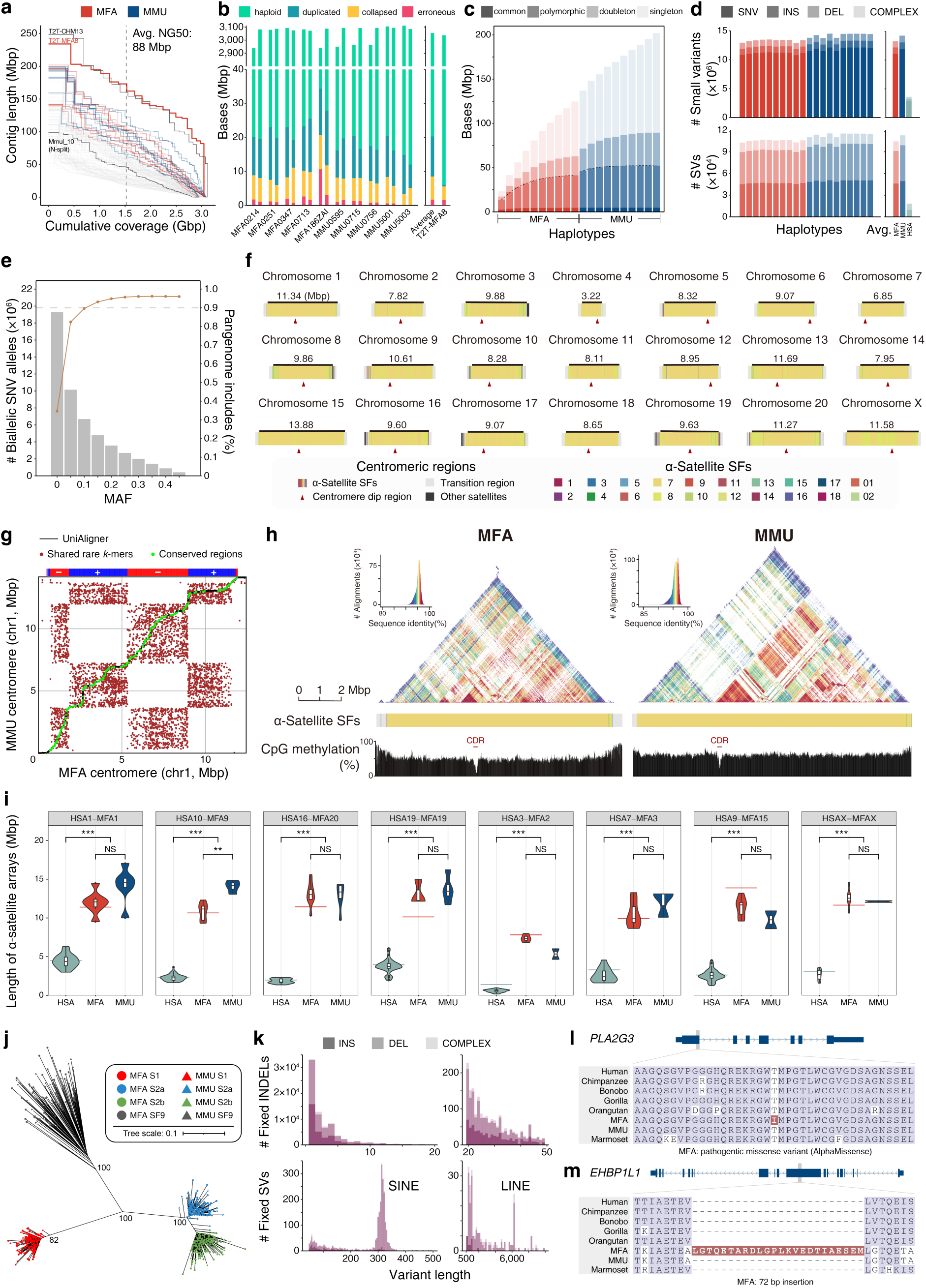
20 haplotype-resolved macaque assemblies, pangenome graph, centromere, and lineage-specific genetic variation. **(a)** The cumulative distribution of genome length shows 10 haplotype-resolved MFA assemblies and 10 MMU assemblies in red and blue, respectively. T2T-MFA8, T2T-CHM13, and Mmul_10 (split by Ns) are labeled for comparison. The 94 human genome assemblies in HPRC are represented in light gray. The average NG50 of macaque assemblies is 88 Mbp, suggesting the high contiguity of macaque assemblies. **(b)** Flagger evaluation of 20 haplotype-resolved assemblies is shown on the left panel, while the average of 20 assemblies and T2T-MFA8 is shown on the right panel. **(c)** The cumulative number of added bases when adding assemblies one by one is illustrated, with red representing MFA and blue representing MMU. The total of added sequences shows slow growth after the seventh MFA or MMU assembly. The species switch (MFA→MMU) increases the yield of added sequences. Transparent colors indicate singleton (AF < 5%), doubleton (5% ≤ AF < 10%), polymorphic (10% ≤ AF < 50%), and common (AF ≥ 50%) alleles. **(d)** The number of genetic variants in each haplotype. The left panel shows the number of small variants (top) and SVs (bottom) per haplotype in the pangenome graph. The right panel shows the average number of small variants (top) and SVs (bottom) of MFA, MMU, and humans (from the HPRC MC pangenome graph). **(e)** The biallelic SNV comparison between the pangenome graph and the macaque whole-genome sequencing (WGS) cohort. The MAF distribution of biallelic SNVs in the largest macaque cohort is displayed in a gray histogram, while the percentage of SNVs in the largest macaque cohort covered by the pangenome graph is shown in a line chart. The panel shows that the pangenome graph covers 80% of genetic variation with MAF ≥ 5% in the largest macaque cohort. **(f)** The complete centromere assemblies of T2T-MFA8v1.0. The different colors refer to the suprachromosomal family (SF) of α-satellite and the length of α-satellite arrays is indicated. The centromere dip regions (CDR) are marked with triangles, as obtained by methylation calling. **(g)** The α-satellite arrays of chr1 dot plot between MFA and MMU generated with UniAligner. The red dots refer to the common rare *k*-mers (*k* ≥ 80) and the green dots refer to the conserved regions between two centromeres. The black line indicates the optimal rare-alignment path. The α-satellite array strand track is shown above the dot plot (blue for forward strand (+) and red for reverse strand (–)). **(h)** Comparison of the MFA and MMU chromosome 1 α-satellite arrays with SF and methylation annotation. The sequence similarity of the 5-kb block is illustrated using StainGlass, with the CDRs indicated by methylation level. **(i)** The length distribution of α-satellite arrays in HSA, MFA, and MMU. The green, red, and blue violin plots represent the length distribution of α-satellite arrays for HSA, MFA, and MMU, respectively. The horizontal lines indicate the length in reference genomes (green for T2T-CHM13 and red for T2T-MFA8). Values in boxes indicate the median and 95% confidence interval. The *P* values are estimated with the Wilcoxon test. NS: not significant, *: *P* < 0.05, **: *P* < 0.01, ***: *P* < 0.001. **(j)** The phylogenetic tree shows that the S1 (red), S2a (blue), S2b (green), and SF9 α-satellites (dark gray) of MFA (round) and MMU (triangle) are clustered into separate clades. **(k)** Lineage-specific fixed genetic variation. The length distribution of fixed INDELs and SVs are shown in the left panel (INDEL: 2-20 bp (top), SV: 50-500 bp (bottom)) and right (INDEL: 20-50 bp (top), SV: 500-10000 bp (bottom)). Notable peaks for *Alu* and *L1* are at 300 bp and 6000 bp. A fixed SNV in *PLA2G3* **(l)** and a fixed SV in *EHBP1L1* **(m)** result in amino acid differences between MFA and MMU.

We next utilized T2T-MFA8v1.0 and the 20 macaque assemblies in a Minigraph-Cactus^71^ (MC) pangenome analysis to characterize the major genetic differences between MFA and MMU. Within this MC pangenome graph, we added 202.31 Mbp of non-reference sequence, of which 47.48 Mbp was classified as polymorphic (present in 3 to 10 haplotypes) and 4.98 Mbp of common genome sequences (present in 10 or more haplotypes) (Figure 3c). The twentieth genome assembly barely contributes to both major and polymorphic genomic sequences of the pangenome, implying that our draft macaque MC pangenome graph already captures a considerable proportion of the major variants within the macaque species (Figure 3c).

Based on the MC pangenome graph, we characterized 50,130,400 SNVs (1 bp), 12,654,525 INDELs (2-49 bp), and 1,063,693 structural variants (SVs; ≥50 bp). A comparison across individual haplotypes revealed that they share similar variant abundance and variant type distribution, with an average of 11.50 million SNVs, 2.27 million INDELs, and 0.11 million SVs per haplotype (Figure 3d). Moreover, the number of various variants in macaques is at least three times higher than in humans, highlighting a greater level of genetic diversity in macaques^31^ (Figure 3d). When using biallelic SNVs as a proxy for estimating the genetic diversity of macaques in the MC pangenome graph, approximately 35% of SNVs identified in the largest macaque cohorts were represented in the MC pangenome graph^11^ (Figure 3e). Moreover, 82% and 90% of SNVs with a minor allele frequency (MAF) exceeding 5% and 10% in the cohorts were included in the MC pangenome graph (Figure 3e), respectively. This suggested that most polymorphic alleles of the macaque genomes (MAF ≥ 5%) are represented in the MC pangenome graph and are suitable for detecting major genetic differences between MFA and MMU in this study. However, it may overlook the rare alleles (MAF < 5%) within macaque populations.

We also characterized genetic variants in the 20 macaques using DeepVariant^72,73^, PBSV^74^, PAV^75^, SVIM^76^, and SVIM-asm^77^ based on their PacBio HiFi reads and haplotype-resolved assemblies (Methods). We successfully identified 61,746,817 high-quality SNVs and INDELs supported by both DeepVariant and PAV (Supplementary Figure 21). Additionally, 442,026 SVs, including 283,690 insertions and 158,336 deletions, were supported by at least one read-based caller (PBSV or SVIM) and one assembly-based caller (PAV or SVIM-asm) (Supplementary Table 17). A comparative analysis between the genetic variants detected in the MC pangenome graph and those identified by the aforementioned callers showed a significant degree of concurrence for SNVs and INDELs with concordance rates of 92.03% and 84.75%, respectively, in euchromatic genomic regions (Supplementary Figure 22). The concordance for SVs, however, was found to be lesser (79.04% for SVs) (Supplementary Figure 22). By excluding the complex genomic regions, including SDs, common repeats, and tandem repeats, we observed an increased level of congruity for SNVs, INDELs, and SVs between the MC pangenome graph and the caller data (SNVs: 95.87%, INDELs: 89.06%, SVs: 94.58%) (Supplementary Figure 22). These findings suggest that the variants identified from the MC pangenome graph are accurate, but current computational tools have limitations in precisely characterizing variants within complex genomic regions.

Additionally, we identified 187 inversions (length: 10 kbp to 4 Mbp) by two independent tools (PAV^75^ and LSGvar) among macaques (Supplementary Figure 23 and Supplementary Table 18). Among 187 inversions, only one was confirmed to be fixed through genotyping the 20 haplotype-resolved macaque genomes (*F_ST_* ≥ 0.8) (Supplementary Table 18). To explore the potential biological implications of these inversions, we examined the expression profiles of genes located in the 500 kbp flanking regions of the inversion breakpoints. Leveraging our previously published RNA-seq data, we found that 43 out of 1,099 protein-coding genes located within the 500 kbp flanking regions of the inversion breakpoints showed differential expression (Supplementary Figure 24 and Supplementary Table 19). However, we did not observe a difference in the proportion of differentially expressed genes (DEGs) relative to the number of genes located within the 500 kbp regions flanking the breakpoints (3.91%) compared to the whole-genome level (3.65%) (chi-square test, *P* = 0.71) (Supplementary Table 20).

### Centromere structure, variation, and diversity within macaques

The centromere is one of the diverse and complex regions in genomes, yet the MC graph excludes centromere regions due to limited alignment algorithms, particularly failing in the long α-satellite arrays that form centromeres. Each primate lineage features active centromeres comprised of distinct α-satellite suprachromosomal families (SFs), and macaque centromeres notably showcase extensive SF7-derived arrays flanked by smaller^18,24,49^, older pieces from SF8-SF13, remnants of ancient centromeres shared with ape lineages^18,24,78,79^ (Supplementary Figure 25). Here, we introduce a new annotation and visualization tool to explore centromere structure, variation, and diversity in the central parts of macaque centromeres (Methods). These regions consist of a homogeneous core formed by S1S2a and S1S2b dimers (∼85.2%), alongside smaller and more divergent layers featuring 3-mers, 6-mers, and X monomers (∼13.2%) potentially from extinct macaque ancestors. The length of α-satellite arrays varies from 3.22 Mbp (chr4) to 13.88 Mbp (chr15) in T2T-MFA8v1.0, with sequence similarity in the 5 kbp continuous blocks ranging from 71.4% to 100.0% (Figure 3f and Supplementary Figure 26).

Next, we investigated whether macaque α-satellite arrays vary between MFA and MMU. Comparing the S1S2 α-satellite arrays in our 20 haplotype-resolved genome assemblies, 139 of 420 centromeric regions (33.10%) were successfully assembled, with lengths varying from 2.56 to 17.05 Mbp (Supplementary Table 21). Both MFA and MMU α-satellite arrays, similar to apes, exhibit sequence similarity block structures with distinct centromere dip regions^79^ (Figure 3g, h and Supplementary Figure 26; Great ape T2T assemblies, in prep). Additionally, the α-satellite arrays of chromosome 1 are conserved between MFA and MMU, with the ancestral macaque-specific centromere layers preserved in both species (Figure 3g and 3h). However, the conservation or synteny degree of α-satellite higher-order repeats between two humans (T2T-CHM13 and T2T-CHM1) varies considerably^24^. Moreover, we found that the lengths of α-satellite arrays between MFA and MMU are not significantly different in most chromosomes, but on average, they are ∼3.7-fold larger than those in humans^18,24^ (Wilcoxon test, *P* = 4.20×10^-12^ for chr1 between MFA and HSA, and *P* = 0.07 for chr1 between MFA and MMU) (Figure 3i). The phylogenetic analysis of 600 randomly chosen α-satellite monomers from five chromosomes reveals that the S1 and S2 α-satellites of MFA and MMU cluster into two distinct clades. Within the S2 α-satellite clade, S2a and S2b split into two separate clades (Figure 3j). In addition, we observed the chromosome-specific S2a, but not for S1 and S2b (Supplementary Figure 27). This chromosome-specific α-satellite is observed for the first time in primate centromeres.

### Genotyping and fixed genetic variation

To gain a deeper insight into both the fixed and polymorphic genetic variants in MFA and MMU, we conducted Illumina short-read WGS on 116 MFA individuals and 35 Chinese rhesus macaque (CMMU) individuals with a mean coverage of 35×. Along with 43 CMMU individuals and 95 Indian rhesus macaque (IMMU) individuals from previously available data^11^, we finally selected 94 unrelated MFA, 67 unrelated CMMU, and 88 unrelated IMMU individuals^80^ (π ≤ 0.25, indicating fourth-degree relatedness, Supplementary Figure 28 and Supplementary Table 22) to genotype 49,558,493 SNVs, 12,537,007 INDELs, and 829,478 SVs in the MC pangenome graph with PanGenie^81^.

As part of our quality control measures, we compared the AFs derived from the 20 haplotype-resolved assemblies in the MC pangenome graph and the AFs generated by PanGenie from this panel of the 249 unrelated macaque individuals. Implementing a machine-learning approach to exclude low-quality genotypes, we curated a high-quality genotype dataset consisting of 46,833,568 SNVs, 9,070,092 INDELs (3,975,746 small insertions, 4,567,595 small deletions, and 526,751 complex INDELs), and 810,024 SVs (596,299 insertions, 138,070 deletions, and 75,655 complex SVs). The correlation of AFs between the MC pangenome graph and PanGenie showed Pearson correlation coefficients of 0.96, 0.94, and 0.88 for SNVs, INDELs, and SVs, respectively (Supplementary Figure 29). This result demonstrated the high quality of the genotype set.

In the genotype set, we identified 558,157 fixed SNVs (∼558 kbp), 76,997 fixed INDELs (∼284 kbp), and 2,797 fixed SVs (∼1,333 kbp). Our data showed that >84% of the fixed-INDEL divergences were predominantly 1-5 bp in size (Figure 3k). Concurrently, 2,791 fixed-SV differences were small (<10 kbp in length) with two notable peaks at 300 bp and 6 kbp indicative of *Alu* and *L1* retrotransposition events (Figure 3k), respectively. In addition, 0.39% of the fixed variants (2,365 SNVs, 112 INDELs, and 15 SVs) lead to 10,165 moderate-impact variants and 156 high-impact variants by Ensembl Variant Effect Predictor^82^. These variants affect 1,503 genes and 6,390 transcripts in macaques (Supplementary Table 23).

These particular genes that intersected with the fixed genetic variants are enriched in cell adhesion (*P* = 4.58×10^-3^), aerobic respiration (*P* = 1.95×10^-3^), and DNA repair^67^ (*P* = 6.46×10^-3^) (Supplementary Table 24). Of special interest, a 1 bp fixed SNV distinguishes MFA from MMU and other apes within *PLA2G3*, which encodes a secretory calcium-dependent phospholipase A2 protein involved in mast cell maturation^83,84^ (Figure 3l). Similarly, a fixed 72 bp insertion in *EHBP1L1* in MFA, with respect to MMU and other apes, led to a 24 amino acid insertion. *EHBP1L1* is involved in apicobasal polarity and cilia length regulation^85,86^ (Figure 3m). To further validate our findings, we confirmed these genetic variants using Iso-Seq data (Supplementary Figure 30). The *PLA2G3* gene encodes an enzyme involved in lipid metabolism, and its knockout in mice leads to defects in sperm maturation and fertility^87^. *EHBP1L1* is involved in the Rab8/10-EHBP1L1-Bin1-dynamin signal transduction pathway, which is essential for maintaining cell polarity^85,86^. Loss of *EHBP1L1* results in lethal anemia in mice^85^.

### Gene copy number variation within macaque populations

We identified 370 SDR hotspots (∼240 Mbp) based on the bubbles greater than 10 kbp in the MC pangenome graph (Supplementary Table 25, Methods). Within these hotspots, we discovered 17 genes, including *Mafa-AG/B*, *CYPs*, *GSTMs*, and *RNASEs*, each comprising at least four copies (Figure 4a, Methods). Meanwhile, we genotyped the copy number (CN) of each gene in the 249 unrelated macaque individuals. To quantify the extent of gene CN variation, we implemented the Shannon diversity index and identified 26 genes that showed significant variability between MFA and MMU (Shannon diversity index ≥ 1) (Figure 4b).

**Figure 4.**
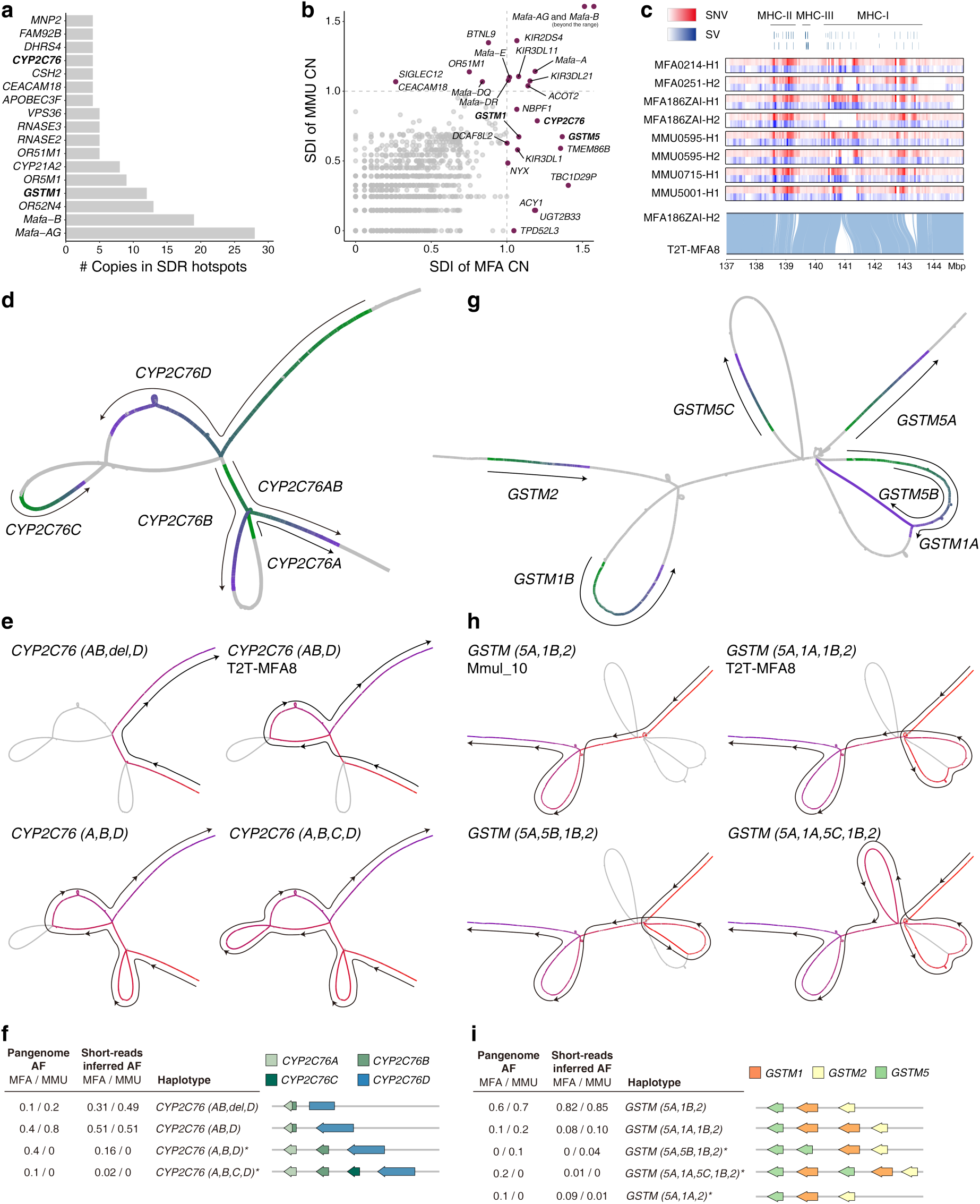
Genic copy number variation in complex loci. **(a)** The most common copy number (CN) variable genes in SDR hotspots of macaques. The x-axis represents the number of gene copies that can be mapped to a bubble in the pangenome graph, while the y-axis shows the 17 most CN variable genes. **(b)** Comparison of CN differentiation between MFA and MMU. The diversity degree of CN variation in MFA and MMU is represented with the Shannon diversity index (SDI) on the x-axis and y-axis, respectively. An SDI ≥ 1 is used to determine the gene CN diversity. **(c)** The complexity of major histocompatibility complex (MHC) in macaques. SNV and SV densities for eight structural haplotypes with gene models are shown above (top). The syntenic relationship between T2T-MFA8 and MFA186ZAI-H2 (bottom) shows a ∼1 Mbp deletion in MFA186ZAI-H2 with respect to T2T-MFA8. **(d)** The structural haplotypes of *CYP2C76* copies. Green and purple refer the start and end of *CYP2C76* gene bodies, respectively. **(e)** The graphical representation of four structural haplotypes of *CYP2C76* follows different paths in the pangenome, with red and purple representing the start and end of a path, respectively. The haplotype of T2T-MFA8 is *CYP2C76 (AB, D)*, while this region is unresolved in Mmul_10. **(f)** Frequency statistics of *CYP2C76* haplotypes and their schematic graph. The frequency of structural haplotypes in the pangenome graph is displayed in the first column, while the inferred frequency from the population with short-read genotyping is shown in the second column. **(g-i)** The structural haplotype of the *GSTM* gene family in the pangenome graph, with visualization presented in the same format as (d-f).

We next analyzed the genes that both were contained in the SDR hotspots and exhibited CN differences between MFA and MMU. *Mafa-AG* and *Mafa-B* (*HLA-G* and *HLA-B* in humans) in the major histocompatibility complex (MHC) Class I region showed the largest CN diversity between MFA and MMU (Figure 4b). The MHC genomic region is well established for its expansion and diversification in macaque genomes compared to human genomes^10,26,88,89^ (average length in humans: 4.49 Mbp, in macaques: 5.96 Mbp; Wilcoxon test, *P* = 3.5×10^-12^) (Supplementary Figure 31). Here, we identified 12 distinct structural haplotypes (length: 5.44 to 6.65 Mbp) including 269,282 SNVs, 47,129 INDELs, and 6,177 SVs in the MHC region (Figure 4c). Of these genetic variations, the length of 19 single insertion or deletion events is greater than 100 kbp. For example, we found a 407 kbp insertion and a 674 kbp deletion in the maternal haplotype of MFA186ZAI, with respect to T2T-MFA8v1.0, resulting in gain or depletion of 37 genes.

We found that five CN-diverse genes are related to metabolism function (e.g., *CYP2C76*, *GSTM1*, and *GSTM5*). Of particular interest, *CYP2C76* is a monkey-specific gene lacking in apes^79,90,91^ (Supplementary Figure 32; Great ape T2T assemblies, in prep). We newly identified four structural haplotypes (length: 21 to 186 kbp) in macaques, wherein the CN of *CYP2C76* varies from 2 to 4. Of these, one haplotype with *CYP2C76A*, *CYP2C76B*, and *CYP2C76D*, and another with *CYP2C76A*, *CYP2C76B*, *CYP2C76C*, and *CYP2C76D*, are both present in MMU but not in MFA (with pangenome AF of 40% and 10%) (Figure 4d-f). In addition, the *GSTM* gene cluster including *GSTM1*, *GSTM2*, and *GSTM5* showed five structural haplotypes in the MC pangenome graph (length: 40 to 87 kbp). One major haplotype was shared by MFA and MMU, which includes *GSTM1B*, *GSTM2*, and *GSTM5A*. The other four minor haplotypes showed a slight discrepancy between the two species (Figure 4g-i). We characterized the AFs of these haplotypes not only within the MC pangenome graph but also inferred their AFs through genotyping the aforementioned panel of 249 unrelated individual macaques (Figure 4f and 4i, Supplementary Tables 26 and 27) (Methods). The major haplotypes identified in the *CYP2C76* and *GSTM* regions through both the MC pangenome graph and short-read genotyping showed concurrence. Yet, there was a modest degree of variation in the AFs of minor haplotypes. One possibility is that our MC pangenome graph is capable of inferring the major alleles but struggles to accurately depict the diversity of minor alleles in complex loci.

### Large-scale genomic rearrangement between macaques and humans

Characterizing large-scale genomic rearrangements between humans and macaques can enhance research in primate evolution, human genetics, and evolutionary medicine. Previous studies have characterized chromosomal rearrangements between humans and macaques using cytogenetic and other approaches. However, large-scale genomic rearrangements at single-base resolution between humans and macaques remain unexplored. Here, we developed an alignment cluster-based approach (LSGvar, Methods) and identified 159 syntenic blocks between T2T-CHM13v2.0 and T2T-MFA8v1.0 (>100 kbp) with manual curation (Supplementary Table 28). These syntenic blocks are separated by 11 centromere repositions, 76 inversions, and 4 intrachromosomal translocations in primate evolution (Figure 5a, Supplementary Figures 33 and 34, and Supplementary Tables 29 and 30). Among these, 11 centromere repositions, 65 inversions, and 4 intrachromosomal translocations represent fixed genomic differences between humans and macaques (Figure 5a and Supplementary Tables 29 and 30). Notably, 19 large-scale genomic rearrangements (1 centromere reposition, 14 inversions and 4 translocations) are reported for the first time^47,92,93^ (Supplementary Tables 29 and 30).

**Figure 5.**
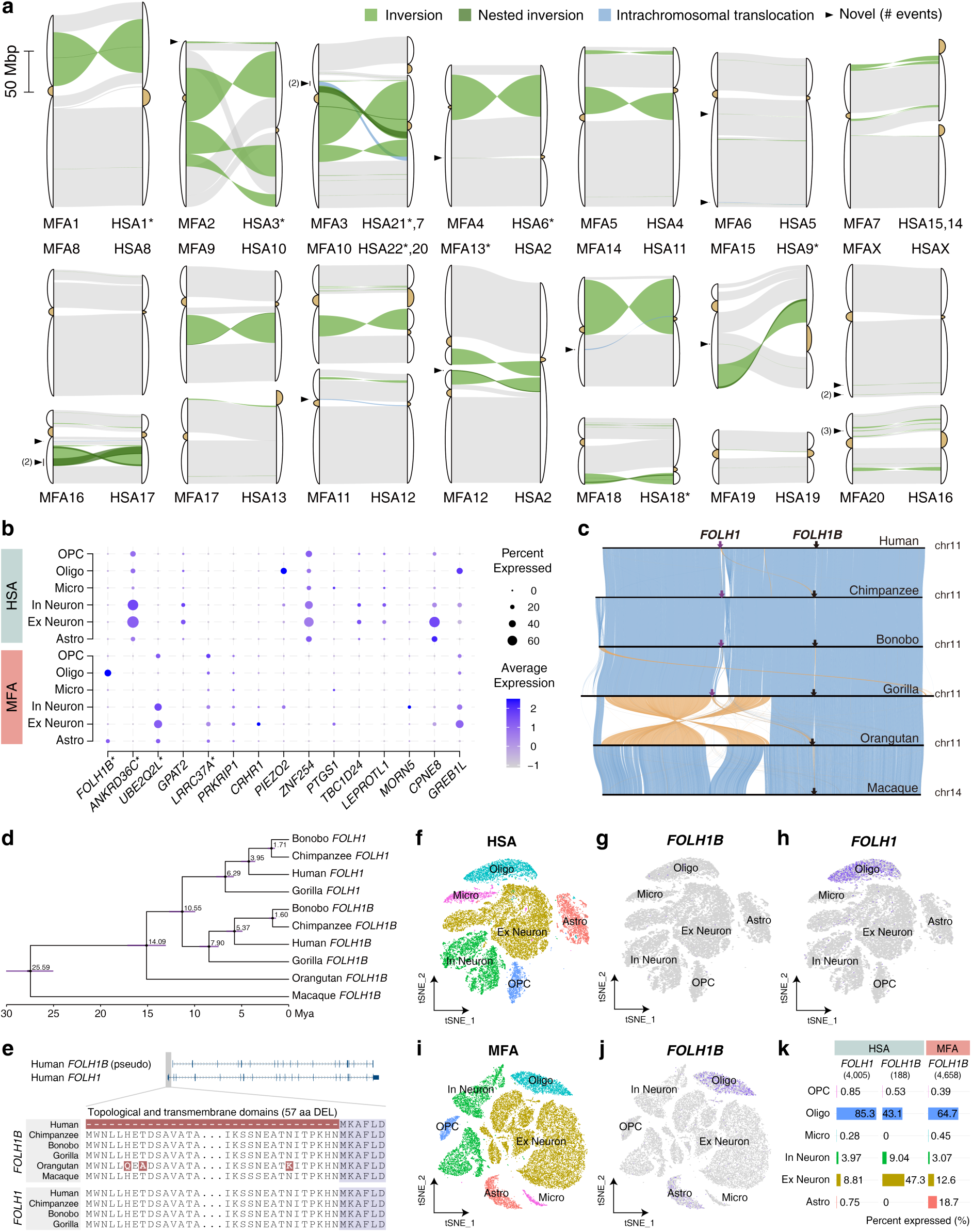
Genomic and gene expression differences between humans and macaques. **(a)** Chromosomal rearrangements between T2T-MFA8 and T2T-CHM13. Macaque and human chromosomes are listed on the left and right for each comparison, respectively. (Inversions in green, and nested inversions in dark green, and intrachromosomal translocations in blue). The 19 newly identified rearrangements are marked with triangles. The asterisk (*) refers to the inverted strand of a chromosome (q arm to p arm). **(b)** Percentage and expression of genes expressed in different cellular types of the prefrontal cortex in humans (HSA) and macaques (MFA). Differential genes located within the 500 kbp flanking regions of large-scale genomic rearrangements are shown on the x-axis, while genes within SDs in primate evolution are marked with an asterisk. Cell types are identified from single-nuclei RNA-seq by specific markers (OPC: oligodendrocyte precursor cells, Oligo: oligodendrocytes, Micro: microglia, In Neuron: inhibitory neurons, Ex Neuron: excitatory neurons, Astro: astrocytes). **(c)** Syntenic relationship of T2T-MFA8 chr14 (apes chr11) in primates and the inversions are shown in orange. The macaque *FOLH1B* (black arrows) is an ancestral copy of human *FOLH1* (purple arrows) and *FOLH1B*. **(d)** The phylogenetic tree of *FOLH1B* and *FOLH1* in primates shows that *FOLH1* and *FOLH1B* are duplicated in the great ape ancestor 10.55 million years ago. The estimated divergence time (above node) with a 95% confidence interval (CI, horizontal purple bar) is calculated using BEAST2 and all nodes have 100% posterior possibility support. **(e)** Amino acid alignment of *FOLH1B* and *FOLH1* in primates shows a fixed human-specific 57 amino acid deletion in *FOLH1B*, which results in the loss of the topological and transmembrane domains and makes *FOLH1B* a pseudogene in humans. **(f-j)** t-SNE visualization with cellular types labeled. Human *FOLH1* (h) is mainly expressed in Oligo, whereas macaque *FOLH1B* (j) is mainly expressed in Oligo and Astro, indicative of differences in gene expression due to the genomic rearrangements. **(k)** The proportion of each gene expressed in each cellular type, with the total expressed cell count shown in brackets.

A total of 2,045 genes are included in the 91 large-scale genomic rearrangements with 500 kbp flanking regions of breakpoints. Among these, the expression of 497 genes is significantly different in the brains of humans and macaques^65^ (Supplementary Figure 35). Notably, 233 genes are upregulated in humans, including *EIF2AK1*, *FAM220A*, and *PIEZO2*, while 264 genes are upregulated in macaques, including *CLN3*, *FOLH1B*, and *APCDD1* (Supplementary Table 31). In addition to the DEG analysis on tissue bulk level, here, incorporating the single-cell RNA-seq data of the dorsolateral prefrontal cortex in humans^94^ and macaques^95^, we identified 444 genes that exhibit both different expression levels in brain tissues and different expression patterns in cell types, including *FOLH1B*, *APCDD1*, and *PIEZO2* (Figure 5b and Supplementary Table 32). Interestingly, 36 of 444 genes are duplicated genes in primate evolution.

Among these genes, *FOLH1* and *FOLH1B* are of particular interest, which typically encode the glutamate carboxypeptidase II (GCPII) involved in the glutamate regulation in neural activity^96^. Mutations in these genes were linked to intellectual disability^97–99^. Our genomic syntenic and phylogenomic analyses showed that *FOLH1* and *FOLH1B* were duplicated in the ancestor of African great apes at 10.55 Mya (95% CI: 9.30-11.81 Mya), and *FOLH1B* is the ancestral orthologous gene in African great apes and macaques^79^ (Figure 5c, d and Supplementary Figures 36 and 37; Great ape T2T assemblies, in prep). Additionally, *FOLH1* and *FOLH1B* contain the same ORF in African great apes, except in humans, where a human-specific deletion alters the ORF (Figure 5e). The 57-amino-acid depletion in human *FOLH1B* represents a fixed mutation in human populations^26^ (AF = 1, HPRC), which results in the loss of the transmembrane domain and is potentially associated with pseudogenization.

Moreover, we found that human *FOLH1* showed high and specific expression in oligodendrocytes (∼85.3% of 4,005 cells), while human *FOLH1B* exhibited minimal expression in brain cells (observed in only 188 cells) (Figure 5f-h). In contrast, macaque *FOLH1B* was expressed in oligodendrocytes (∼64.7% of 4,658 cells), astrocytes (∼18.7% of 4,658 cells), and excitatory neurons (∼12.6% of 4,658 cells) (Figure 5i, j and Supplementary Table 33). These findings suggested significant alterations in the expression patterns of *FOLH1* and *FOLH1B* across different brain cell types between macaques and humans (chi-square test, *P* < 2.2×10^-16^; Figure 5k and Supplementary Table 33). In summary, the evolutionary history and gene expression analysis of *FOLH1* and *FOLH1B* reveal how genomic rearrangements, duplications, and genetic variations alter the gene structure and its regulation in primate evolution.

Another interesting observation is a ∼1.28 Mbp fixed inversion identified between humans and macaques, with the genes *APCDD1* and *PIEZO2* positioned at the 500 kbp flanking region of the breakpoints^47^ (Supplementary Figure 38). We observed that *APCDD1* and *PIEZO2* both exhibited differential expression levels (*P* = 1.54×10^-3^ for *APCDD1*; *P* = 2.61×10^-5^ for *PIEZO2*) and differential cell type patterns (chi-square test, *P* = 5.71×10^-273^ for *APCDD1*; chi-square test, *P* = 0 for *PIEZO2*) between humans and macaques (Supplementary Figure 39). Specifically, in macaques, *APCDD1* was predominantly expressed in excitatory neurons (34.61%), while in humans, it was predominantly expressed in astrocytes (35.96%). Additionally, *PIEZO2* was predominantly expressed in oligodendrocytes in humans (46.19%) but barely expressed in oligodendrocytes in macaques (0.01%). These findings suggest that an inversion event could not only alter gene expression levels but also influence the cell types where the genes are expressed. *APCDD1* is primarily implicated in regulating cell proliferation, differentiation, and development in the nervous system^100^. *PIEZO2* serves as a crucial component of mechanotransduction machinery, responsible for converting mechanical stimuli into electrical signals specifically within brain cells^101,102^. These genes hold potential for influencing brain size and function in primate evolution.

### Population history, speciation, and selection

To deepen our understanding of the population history and selection dynamics between MFA and MMU, we used 26,722,181 pruned bi-allelic SNVs to perform a principal component analysis (PCA). As expected, the PCA results depicted a clear grouping of MFA, CMMU, and IMMU into distinct clusters (Figure 6a). We also investigated the long-term demographic history of MFA, CMMU, and IMMU with the multiple sequentially Markovian coalescent (MSMC) approach^103^. Our findings showed a population contraction within the macaque species during the Pliocene (2.5-5.3 Mya) (Figure 6b). Following this period, the effective population size (*N_e_*) of CMMU and IMMU showed a steady increase, while *N_e_* of MFA peaked at ∼0.6 Mya (Figure 6b).

**Figure 6.**
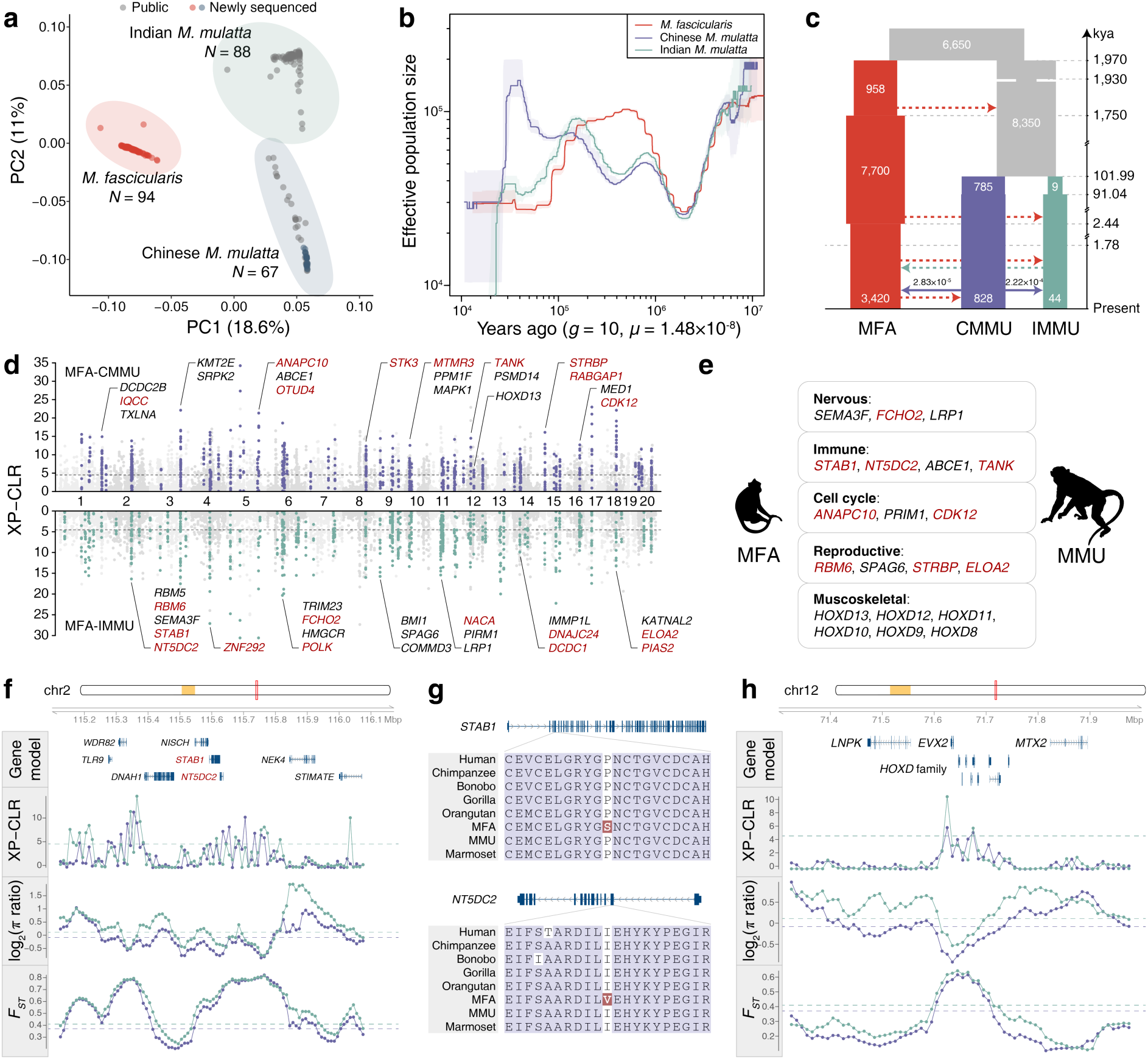
Demographic history and selection. **(a)** Principal component analysis (PCA) of three macaque populations. The first component (18.6%, x-axis) separates MFA (red) and MMU, while the second component (11%, y-axis) distinguishes CMMU (purple, Chinese rhesus macaque) and IMMU (green, Indian rhesus macaque). The macaque individuals are clustered according to each population. Newly sequenced samples in this study are marked in color, while the samples from previous study are marked in gray. **(b)** Demographic history inference of 10 long-read macaque genomes using MSMC2. Effective population size (*N_e_*) is estimated from heterozygosity, with a generation time (*g* = 10) and an estimated mutation rate per generation (*µ* = 1.48×10^-8^). The median size for each population group is shown by the line, and the area between the upper and lower quartiles is shaded. **(c)** Speciation with gene flow model of MFA, CMMU, and IMMU. The population split time, *N_e_*, and migration rate are estimated from a site frequency spectrum using the best-fitting model inferred by fastsimcoal2. The graphic summary shows estimated migration rates (less than 1×10^-6^ are marked as dashed lines) and detailed demographic history inference can be found in Supplementary Figures 40-42. **(d)** Analysis of selection signals between MFA and MMU. The Manhattan plots (top: MFA vs. CMMU; bottom: MFA vs. IMMU) illustrate the normalized XP-CLR score (y-axis). The horizontal gray lines represent the top 1% threshold from XP-CLR. Selected loci between the two populations (top 1% XP-CLR, top or bottom 5% *π* ratio, and top 10% *F_ST_*) are marked in purple or green. Genes with fixed amino acid changes are marked as deep red. **(e)** The functional enrichment of genes under selection. The genes under selection are enriched in neurodevelopment, immunity, cell cycle, reproduction, and skeletal development. **(f)** A Genomic region associated with *STAB1* and *NT5DC2* shows an obvious selection signal. The gene models, XP-CLR, log_2_(*π* ratio), and *F_ST_* across the genomic region are shown from top to bottom. The dashed lines indicate the top 1% threshold from XP-CLR, the bottom 5% threshold from *π* ratio, and the top 10% threshold from *F_ST_*, respectively. **(g)** Fixed missense variants of *STAB1* and *NT5DC2* result in amino acid differences between MFA and MMU. **(h)** A genomic region associated with the *HOXD* family shows an obvious selection signal.

Next, we used a composite maximum likelihood inference based on site frequency spectrum to infer recent demographic history and gene flow during the speciation and lineage divergence of the three macaque groups. Through testing various isolation and migration models, we found that the model involving weak ancestral and strong recent gene flows significantly fitted the data in fastsimcoal2^104^ (Supplementary Figures 40 and 41, Supplementary Tables 34 and 35, Methods). According to this model, MFA and MMU diverged 1.97 Mya (95% CI: 1.94-1.98 Mya), while the two lineages (CMMU and IMMU) within MMU diverged approximately 101.99 thousand years ago (kya) (95% CI: 101.16-106.98 kya) (Figure 6c and Supplementary Table 36). We also observed two major recent gene flows from CMMU to IMMU (2.22×10^-4^ per site per generation) and MFA (2.83×10^-5^ per site per generation) in the past three thousand years (Figure 6c and Supplementary Figure 42). The *N_e_* changes inferred from the model agreed with the MSMC analysis (Figure 6b and 6c). These results align with the current geographic distribution of macaques, highlighting the widespread distribution of CMMU.

Additionally, we inferred the selective forces that may have shaped macaque speciation. We leveraged an integrated approach (*π* diversity^105^, *F_ST_*^106^, and XP-CLR^107^) to identify the candidate genes involved in the speciation between MFA and MMU. We identified 340 and 326 candidate genes associated with selective sweeps during the speciation of MFA and CMMU, and MFA and IMMU, respectively (Figure 6d and Supplementary Table 37); 146 genes appeared to be common to both MFA-CMMU speciation and MFA-IMMU speciation (Figure 6d). These genes are also implicated in various biological functions, including neural development (e.g., *SEMA3F* and *LRP1*), immune response (e.g., *STAB1* and *NT5DC2*), cell cycle progression (e.g., *ANAPC10* and *CDK12*), reproduction (*RBM6* and *ELOA2*), and skeletal development (e.g., *HOXD* gene cluster) (Figure 6e and Supplementary Table 38).

Among the 146 genes, 36 (24.66%) showed fixed amino acid differences between MFA and MMU. For example, MFA-specific amino acid changes are observed in STAB1 (Pro→Ser) and NT5DC2 (Ile→Val) compared to other primates (Figure 6f and 6g). *STAB1* encodes a transmembrane receptor protein involved in lymphocyte homing or receptor scavenging^108,109^, while *NT5DC2* belongs to the family of haloacid dehalogenase-type phosphatases^110^. Both *STAB1* and *NT5DC2* are known to be associated with bipolar disorder and schizophrenia^111–113^. Immune dysfunction is a major factor in the pathophysiology of bipolar disorder and other psychiatric diseases^114^. Yet, the specific functional implications of these amino acid changes between MFA and MMU in immune and neuronal processes remain unclear.

Of the 146 genes, 110 (75.34%) did not show fixed amino acid changes between MFA and MMU (e.g., *SEMA3F* and *HOXD13*). *SEMA3F* plays a crucial role in gonadotropin-releasing hormone neuron development, and its loss of function leads to idiopathic hypogonadotropic hypogonadism in humans^115,116^. *HOXD13*, a member of a highly conserved family of transcription factors, is essential in mammalian morphogenesis^117–119^. The recent expression quantitative trait loci analysis in deer mice suggests that the expression difference of *HOXD13* is a significant determinant in tail length variation for environmental adaptation^120^. In our study, we also observed a strong selection on the *HOXD13* locus when comparing MFA and MMU. Given the longer tail in MFA compared to MMU, we hypothesize that *HOXD13* plays a similar role in macaque tail development. If confirmed, this would represent another example of convergent evolution. In summary, these findings suggest a potential association between genes related with selective sweeps and the observable phenotypic variations (e.g., tail length, reproduction, and others) between MFA and MMU (Figure 6h). However, to substantiate these associations, additional functional assays are necessary.

## DISCUSSION

Macaques play crucial roles in biomedical research and the study of primate evolution^11,27,28,47^. In this study, to enhance our understanding of macaque genome structure, population history, major genetic differences between the two important macaque species, and their comparison with modern humans, we generated the parthenogenetic cell line, provided abundant data, and performed integrated analyses. Overall, these combined efforts can substantially advance research in the NHP fields regarding biomedical model selection, gene editing, gene regulation, and primate evolution. Additionally, our refined analyses of selection and population history within the two macaque species offer novel insights into macaque speciation and their phenotypic adaptation.

The first T2T macaque genome assembly (T2T-MFA8v1.0) facilitates the identification of the genome structure differences between macaques and humans^19,55,56^. First, SD differences exist between the macaque and human genomes in various aspects such as their length, the proportion of intra and interchromosomal SDs, and their genomic position (Supplementary Figures 12 and 13). Second, the rDNA array in macaques is only found in chromosome 10, in contrast to humans, where they are located in five chromosomes^15,121^ (chr13, chr14, chr15, chr21, and chr22) (Figure 1e). Third, the length and composition of the α-satellite arrays differ between macaques and apes^18,24,49^. Additionally, while the α-satellite arrays are relatively conserved within macaques, they exhibit greater variability in humans^18,24^. Finally, 91 large-scale genomic differences between macaques and humans have been revealed at single-base resolution, most of which are associated with centromere repositions, inversion, and translocation^47,92,93^ (Figure 5). The detailed analysis of *FOLH1* and *FOLH1B* shows the complicated fate of duplicated genes and the gene regulation changes caused by large-scale genomic structural divergence. In conjunction with the case of *APCDD1* and *PEIZO2*, our study proposes a concept that these genomic differences influence not only the gene expression level but also the cellular expression patterns of genes. These efforts contribute to our understanding of gene functional alteration by large-scale rearrangements between humans and macaques, which is crucial for understanding lineage-specific phenotype^11,33–35^ (e.g., cortex expansion) and human diseases.

The MC pangenome graph, which includes 20 haplotype-resolved macaque genomes, reveals a higher genetic diversity in macaques compared to humans^26^. We have identified 2.17 Mbp of fixed genetic variation between MFA and MMU. These genetic variations are associated with 1,503 protein-coding genes enriched in immune response, tail development, and metabolism (Supplementary Table 24). We also identified 240 Mbp complex loci in MFA and MMU and discovered the haplotype diversity of *CYP2C76* and *GSTM* gene family with the pangenome. In addition, with the advanced full-length RNA deep-sequencing technologies, the analysis of alternative splicing differences provides new insights into the genetic differences at the transcriptome level between MFA and MMU. These major genetic differences provide a roadmap for further understanding the functional biological differences of the two macaques, which is important for future biomedical research^60–64^.

Previous studies have shown gene flow between MFA and MMU. Here, our population genomics analysis provides a more detailed history of speciation with gene flow in macaques^103,104^. By leveraging data from 249 unrelated macaque individuals, we identify 146 genes with selective sweeps in these two species, and these genes are enriched in various biological functions associated with adaptation (Supplementary Tables 37 and 38). Intriguingly, when integrated with our macaque fixed-genetic-difference atlas, we found that only 24.66% of genes in the regions with selective sweeps contain variants resulting in amino acid changes (chi-square test, *P* = 1.52×10^-14^) (Supplementary Table 39). This suggests that the majority of phenotypic differences may arise from gene regulation rather than alterations in protein sequences, similar to the case of *HOXD13* in deer mice tail length^120^. Nonetheless, functional assessments of these candidate genes are needed to investigate the phenotypic differences between these two macaques.

In summary, this study both establishes a robust genetic foundation for the macaque biomedical models and also improves our understanding of primate evolution, human diseases, and lineage-specific adaptation. However, there are several limitations. First, the present study primarily focuses on characterizing major allele differences between species, potentially overlooking rare and minor alleles. Second, since all macaque individuals are sourced from primate research facilities in China and the USA, the study may not fully capture the natural genetic diversity and population history of wild macaques. Therefore, collecting additional bio-samples from wild macaques using noninvasive approaches and performing more extensive long-read macaque genome sequencing are warranted. Finally, while we highlight gene regulatory alterations stemming from large-scale genomic divergences between humans and macaques, there remains a deficiency of gene regulation data across various developmental stages, akin to an ENCODE dataset for macaques. Such data are crucial for fully elucidating the functional consequences associated with these genetic variations.

## Methods summary

We generated a parthenogenesis cell line, MFA582-1, from an MFA, from which we produced 53× HiFi, 77× ONT, 32× Illumina WGS, and 156× Hi-C data. Using these data, we assembled a complete genome (T2T-MFA8v1.0) using Hifiasm^21,23^, Verkko^22^, and our SUNK-based pipeline. The genome assembly was annotated with 15.27 million FLNC reads (Iso-Seq) of MFA across 15 different tissues, including 6 brain tissues. 14.82 million FLNC reads from 15 tissues of MMU were also sequenced, and we used these data to perform a comparative transcriptome analysis with IsoSeq3 and SQANTI3^122^. SD analysis using SEDEF^123^ and protein structure prediction using AlphaFold2^66^ were performed. We sequenced 10 macaque individuals using HiFi and ONT and assembled their genomes with Hifiasm^21,23^. A draft pangenome graph was generated using MC^71^. Genetic variations, including SNVs, INDELs, and SVs, were called by graph decomposition. Another genetic variation callset was generated from the callers with DeepVariant^72,73^, PBSV^74^, PAV^75^, SVIM^76^, and SVIM-asm^77^. PanGenie^81^ and fastCN^124^ were used to genotype the SVs and copy number variants from a diversity panel of 249 unrelated macaque individuals. A cluster-based approach (LSGvar) was developed to characterize large-scale rearrangements between macaques and humans. These large-scale genomic differences were genotyped using 94 haplotype-resolved human^26^ (HPRC) and 20 macaque genomes. We used DESeq2^125^, Limma^126^, Cell Ranger^127^, and Seurat^128,129^ to investigate gene expression and cell type differentiation of genes associated with these large genomic difference regions. Demographic modeling was conducted using MSMC2^103^ and fastsimcoal2^104^, and selection signals were examined using three independent approaches: *π* diversity^105^, *F_ST_*^106^, and XP-CLR^107^. More detailed information can be found in the Supplementary Information.

## Ethics statement

The usage of research animals in this study underwent evaluation and approval by the Primate Life Sciences Ethics Committee of the Center for Excellence in Brain Science and Intelligence Technology, Chinese Academy of Sciences (CEBSIT-2021074). In line with reduction principles, surgical procedures were conducted with utmost care to minimize pain and discomfort. Painless euthanasia was carried out following appropriate anesthesia and euthanasia protocols. Before euthanasia via exsanguination, monkeys were administered Zoletil 50 injection at a dose of 25 mg/kg to ensure proper sedation.

## Acknowledgments

We thank the staff of the Non-human Primate Facility of the Center for Excellence in Brain Science and Intelligence Technology for their assistance in animal care. We thank the HPRC and Primate T2T Consortium for providing the long-read human and great ape genome assemblies. We thank Tonia Brown for manuscript proofreading and editing. We thank Kateryna Makova for providing valuable comments. The computations in this study were run on the Siyuan-1 supported by the Center for High Performance Computing at Shanghai Jiao Tong University.

## Funding

This work was supported, in part, by National Natural Science Foundation of China grants (32370658 to Yafei Mao, 82021001 and 31825018 to Q.S., 32300490 to D.W.); by Shanghai Pujiang Program (22PJ1407300) and Shanghai Jiao Tong University 2030 Initiative (WH510363001-7) to Yafei Mao; by the National Key Research and Development Program of China (2022YFF0710901), the Biological Resources Program of the Chinese Academy of Sciences (KFJ-BRP-005), and the National Science and Technology Major Project (Innovation 2030 2021ZD0200900) to Q.S.; by the China Postdoctoral Science Foundation grant (2022M713072 to Y.F.); by National Institutes of Health (NIH) grants (HG002385 and HG010169 to E.E.E., GM147352 to G.A.L., and R01HG011274-01 to K.H.M.); by the Intramural Research Program of the National Human Genome Research Institute, National Institutes of Health to A.M.P.; by the Center for Integration in Science of the Ministry of Aliyah, Israel to I.A.A.; by PRIN 2022-2022E8NN2N to M.V.; by PRIN 2020-2020J84FAM and PRIN 2022 PNRR-P2022ZE75A to F.A.. E.E.E. is an investigator of the Howard Hughes Medical Institute.

## Author contributions

V.A.S is retired from the Institute of Molecular Genetics. Yafei Mao and Q.S. conceived the project; S.Z., N.X., X.Y., Q.S., and Yafei Mao, generated sequencing data; N.X., Yamei Li, X.B., Yong Lu, L.Z., Yuxiang Mao, and Q.S. contributed the macaque samples, maintained the 582-1 cell line, and performed rDNA validation; S.Z., L.F., X.Y., Y.H., D.M., K.M., C.Y., D.W., G.Z., B.S., Q.S., and Yafei Mao assembled genomes, analyzed the data, and performed quality control analyses; A.M.P., and E.E.E. contributed the nonhuman great ape genomes and analyzed the data; S.Z., L.F., Z.Y., E.E.E, and Yafei Mao performed the performed SV analyses; S.Z., L.F., Z.Y., J.H., and Yafei Mao performed large-scale genomic difference analyses; L.d.G., A.P., F.A., and M.V. performed the FISH analyses; S.Z., X.J., F.R., G.A.L., V.A.S., K.H.M., I.A.A., and Yafei Mao performed centromere analyses; S.Z., L.F., Z.Y., X.J., Yuxiang Mao, Q.S., and Yafei Mao performed single-cell RNA-seq and *FOLH1* gene family and *PIEZO2* analyses; S.Z., Y.F., D.W., and Yafei Mao performed the demographic and selection analyses. L.F., R.H., and Q.L. performed the protein structure prediction analyses; S.Z., L.F., Q.S., and Yafei Mao drafted the manuscript. All authors read and approved the manuscript.

## Competing interests

E.E.E. is a scientific advisory board (SAB) member of Variant Bio, Inc. The other authors declare no competing interests.

## Data and materials availability

The raw Illumina, PacBio HiFi, ONT, and Hi-C data of T2T-MFA8 are deposited in NCBI under BioProject accession number PRJNA1037719. The raw PacBio HiFi, ONT, Hi-C, and assemblies of 10 macaque individuals are deposited in NCBI under BioProject accession number PRJNA1041301. The Iso-Seq data are deposited under NCBI BioProject accession number PRJNA1041301. The Illumina sequences of 189 WGS macaque genomes are deposited in NCBI under BioProject accession number PRJNA1041301. T2T-MFA8 and relative data along with a track hub are available on GitHub (https://github.com/zhang-shilong/T2T-MFA8).

